# IsoSpace: Chemistry-Informed Dimensionality Reduction and Automated Isotope Candidate Detection in Imaging Mass Spectrometry

**DOI:** 10.64898/2025.11.30.691393

**Authors:** Meenakshi, Lukasz G. Migas, Katerina V. Djambazova, Martin Dufresne, Jeffrey M. Spraggins, Raf Van de Plas

## Abstract

Imaging mass spectrometry (IMS) experiments simultaneously map thousands of ion species throughout tissue. Its high-dimensionality makes human interpretation difficult and dimensionality reduction (DR) methods are common to facilitate exploration. Traditional DR methods, such as principal component analysis (PCA), are general approaches, designed to work across application domains. They are usually unaware of, and unable to exploit correlations specific to a particular measurement type. Compression-focused, such ‘blind’ methods often deliver physically impossible latent patterns or make feature combinations that confuse rather than aid human understanding. We introduce a novel chemistry-informed DR method, IsoSpace, with built-in awareness of mass spectrometry-relevant patterns. Although generalizable, IsoSpace substantiates ‘chemistry-informed’ as sensitive to potential isotopic relationships. Like traditional DR methods, IsoSpace groups mass-over-charge (*m/z*)-features to deliver a low-dimensional representation of IMS data. Unlike traditional methods, IsoSpace ensures that retrieved latent patterns constitute potential isotopic families. IsoSpace’s decomposition facilitates IMS interpretation at the molecular species rather than ion species level, implicitly automating isotope candidate detection. Most de-isotoping techniques ignore spatial relationships or rely on molecular class assumptions, making them less suitable for molecularly diverse tissue environments. IsoSpace avoids such assumptions, integrating spatial and spectral cues to empirically detect potential isotopic peaks. IsoSpace uses non-negative matrix factorization and an *m/z* -pattern matrix to uncover isotopic-like sequences, evaluating them by intra-pattern correlation. In a mouse pup example with 879 *m/z* -peaks, IsoSpace identified 71 potential isotopic patterns, substantially reducing data complexity while preserving chemical un-derstanding. IsoSpace o!ers chemical-interpretation-permissive DR with unsupervised isotope candidate detection for heterogeneous samples.

**Figure 0:**
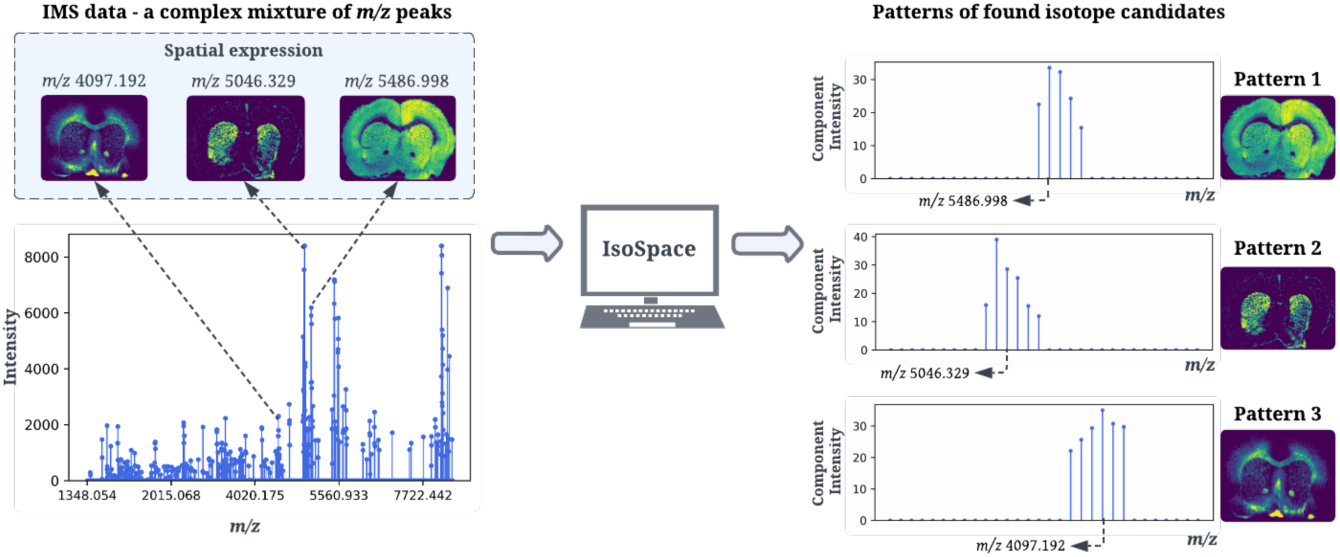
Graphical abstract for IsoSpace, chemistry-informed dimensionality reduction and automated isotope candidate detection in imaging mass spectrometry.

## 1 Introduction

Imaging mass spectrometry (IMS) acquires spatially resolved mass spectra, enabling mapping of chemical compounds throughout biological specimens. ^1–4^ Each spectrum reports hundreds to thousands of ion peaks, offering a comprehensive view into the measurement location’s molecular constituents. However, IMS datasets can become very large and high-dimensional. Therefore, dimensionality reduction (DR) has become essential to IMS analysis, to avoid the “curse of dimensionality” in computational analyses, to reduce data sizes, and to extract meaningful biological patterns underlying these massive datasets. Common IMS-applied DR methods include principal component analysis (PCA),^5^ non-negative matrix factorization (NMF),^4^ t-distributed stochastic neighbor embedding (t-SNE),^6^ and uniform manifold approximation and projection (UMAP).^7^ By capturing dominant variation, these methods are effective for reducing the complexity of high-dimensional data. However, as they are not specifically designed for IMS data, their low-dimensional representations may fail to preserve critical features (*e.g.*, they can dissociate isotopes reporting the same molecular species), which substantially hampers human interpretation of underlying biological pathways.^8,9^

Isotopes are atoms or molecules with the same number of protons but a different number of neutrons. They share nearly identical chemical properties, but differ in mass. Isotopes manifest in IMS data as ion peaks at different *m/z* -positions. Most DR methods are not tailored to detect or preserve these patterns. Isotopic families of peaks can often be viewed as multiple ion species reporting the same molecular species, which can help identify unknown compounds and validate known ones. However, recognizing isotopes in mass spectrometry (MS) measurements can be challenging: different molecular species’ isotopic signatures can overlap and instrumental limitations (e.g., mass resolution, sensitivity) can obscure isotopic features. Therefore, reliable extraction of isotopic information typically requires sophisticated computational algorithms. Several methods to ‘de-isotope’ spectra^8^ have been developed, typically aimed at identifying, removing, or consolidating isotopic peaks to aid the molecular interpretation of a spectrum. Traditional de-isotoping methods focus on analyzing individual spectra,^10^ ignoring IMS-supplied spatial relationships between spectra. Furthermore, many de-isotoping techniques rely on models that assume specific classes of molecules, such as peptides.^11–13^ These models use predefined patterns of isotopic distribution to identify and differentiate between monoisotopic peaks and their isotopic counterparts. ^14^ However, assuming a single molecular class across an entire tissue sample and *m/z* -axis fails to accommodate the true diversity and complexity of biological systems, and limits the applicability of traditional deisotoping methods in IMS’ heterogeneous tissue environments.

Recognizing these challenges, we propose a novel method, IsoSpace, that performs isotope-aware DR for IMS data. Besides a form of chemistry-informed DR, it concurrently facilitates automated detection of potential isotope candidates, without prior assumptions about their molecular class. By integrating spatial and spectral cues, IsoSpace effectively reduces dataset dimensionality, even in heterogeneous, noisy data. It yields a lower-dimensional representation of an IMS dataset that serves the classical purposes of DR, but whose latent axes are also inherently more interpretable for chemists. First, we introduce IMS datasets to evaluate our approach, covering diverse platforms and mass ranges. Then, we present IsoSpace, a two-stage chemistry-informed DR framework. Finally, we demonstrate its application across different IMS case studies.

## 2 Methods and Datasets

Our method is demonstrated on three IMS datasets, covering diverse platforms, tissue types, analyte classes, mass ranges, and pixel sizes. The first dataset is a lipid-focused measurement of a mouse pup whole-body section,^15^ using a matrix-assisted laser desorption/ionization (MALDI) timsTOF flex platform (Bruker Daltonik, Bremen, Germany). The second dataset is a protein/peptide-focused MALDI-IMS measurement of a coronal rat brain section from a Parkinson’s Disease model,^7^ acquired by Fourier-transform ion cyclotron resonance (FTICR) mass spectrometer (Bruker Daltonics, Billerica, MA, USA). The third dataset is a lipid-focused measurement of human kidney tissue from the HuBMAP initiative^1^,^2^, using a MALDI timsTOF fleX (Bruker Daltonik, Bremen, Germany). Figure 1 shows example spectra and ion images for the three datasets, normalized by total ion count (TIC) and peak-picked. Further dataset details, including collection and preprocessing, are available in the supplementary information.

**Figure 1:**
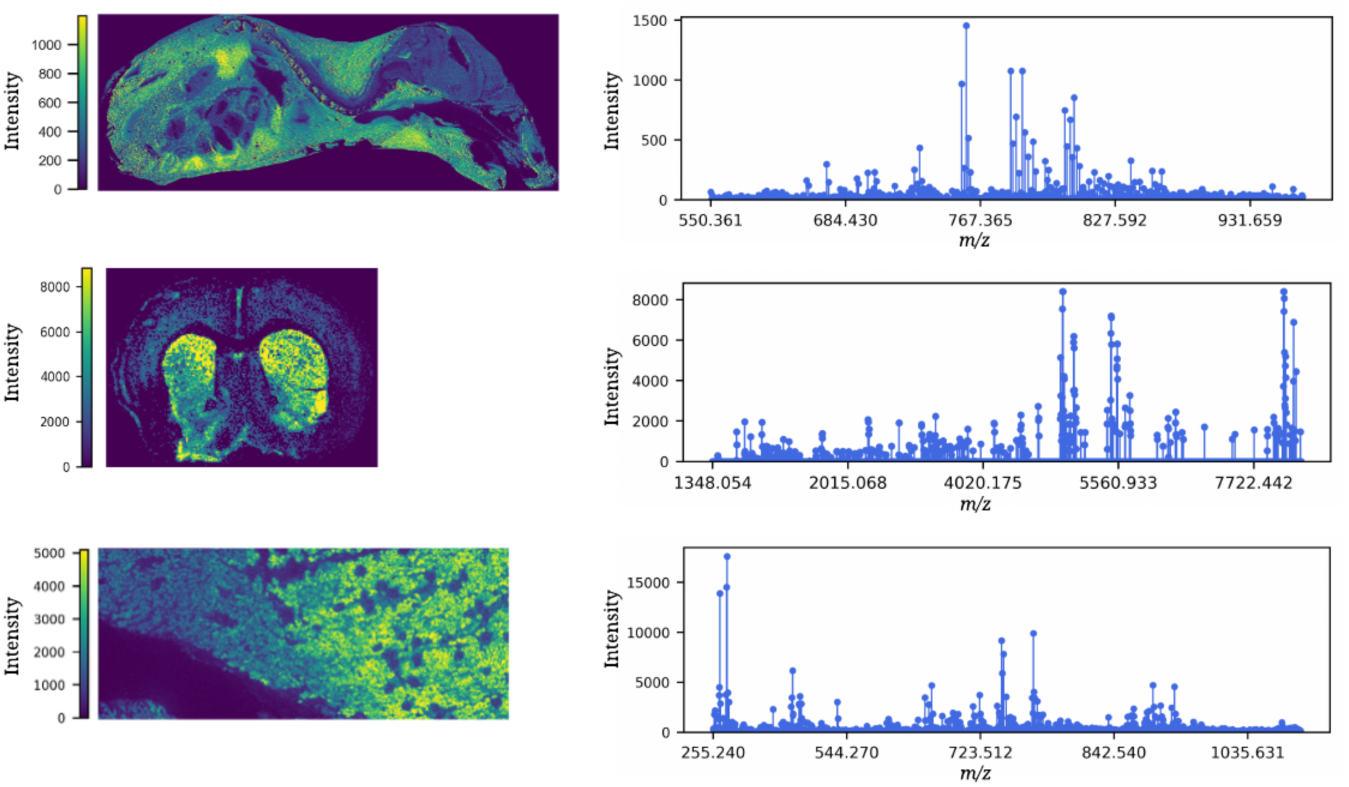
IMS datasets overview. Example ion images and mass spectra for three distinct MALDI-IMS experiments. (top) Whole-body mouse pup section (*m/z* 796.605 image shown). This dataset measures 164,808 pixels at 50-*µ*m pixel size, mapping 879 peaks, and covering a lipid-focused mass range of *m/z* 460-1500. (middle) Coronal section of rat brain (*m/z* 4981.367 image shown). This dataset measures 17,964 pixels at 75-*µ*m pixel size, recording 2611 distinct ion peaks, and covering a protein-focused mass range of *m/z* 1300-23,000. (bottom) Human kidney section (*m/z* 738.508 image shown). This dataset measures 245,744 pixels at 10-*µ*m pixel size, mapping 2199 distinct ion peaks, and covering a lipid-focused mass range of *m/z* 100-2000.

**Figure 2:**
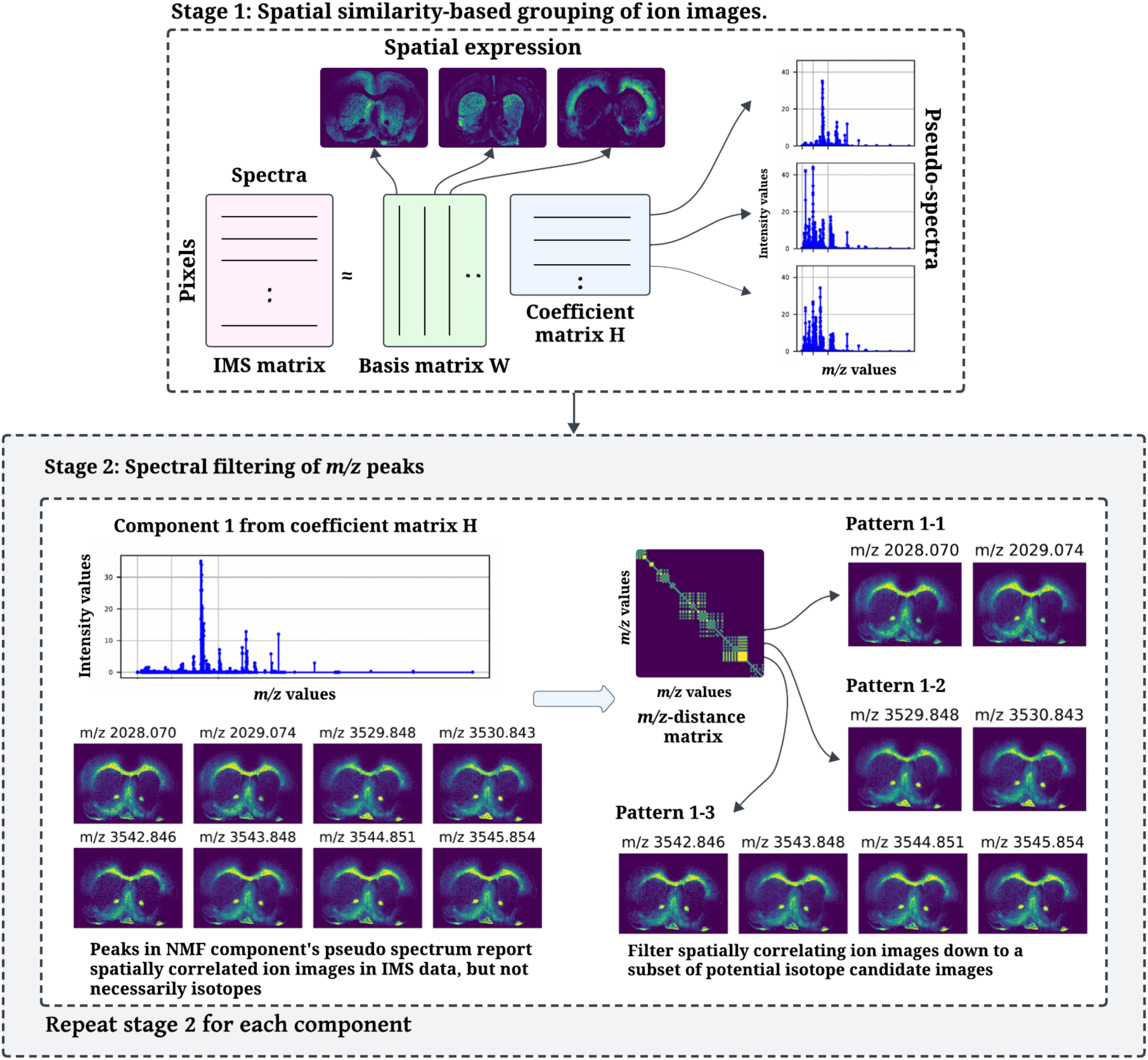
Schematic of the IsoSpace algorithm. (Stage 1) NMF is used to performs dimensionality reduction, grouping ion images that share spatial similarity. (Stage 2) The spatially clustered images undergo further analysis through the lens of the *m/z* -pattern matrix. This matrix is specifically parameterized to uncover potential isotopic patterns along the *m/z* -axis. The NMF-components’ pseudo-spectra are filtered down further into spectral patterns that report potential isotope families.

We now introduce IsoSpace, a two-stage methodology for chemistry-informed DR of IMS data. In this paper, the method is specifically parameterized to detect isotope candidates. However, the algorithm is not tied to isotope-like patterns, and it could be used to search for other *m/z* -distance-based patterns as well. In a first stage, the algorithm analyzes the spatial domain to evaluate spatial coherence between ion distributions and to establish a first shortlist of potential isotope candidates. In the second stage, IsoSpace filters this list for spectral patterns of interest. For this paper, the second stage focuses on *m/z* -proximities that could report an isotopic family. Specifically, two ion peaks are considered isotope candidates of each other if (i) the *m/z* -distance between the peaks fits 1 *± ɛ* (Dalton, if singly-chargedions) with a user-specified uncertainty *ɛ*, and (ii) their ion images exhibit spatially similar tissue distributions, as isotopes would report the same molecular species. This approach yields a lower-dimensional representation of IMS data, but the latent patterns it finds tend to be enriched for isotope candidates, making them interpretable and directly usable by chemists.

### 2.1 IsoSpace stage 1 - Spatial similarity-based grouping of ions

Stage 1 seeks to empirically detect spatial similarity among different *m/z* -peaks’ distributions. It leverages NMF to decompose the IMS experiment’s ion intensity matrix into a reduced set of major NMF-components. Each component groups ion peaks and implicitly enriches for spatial similarity among the ion images it groups. These components empirically reveal which ion images, and thus *m/z* -peaks, spatially correlate.

NMF is widely used to distill IMS data into trends and patterns that make physical sense.^4^ It decomposes an IMS data matrix ***X*** ↓ ℝ*^M↔N^*, where *M* is the number of pixels and *N* denotes the number of spectral bins, into a sum of *k* rank-1 matrices,^16,17^ where *k* ↔ min(*M, N*). The resulting approximation ̃X of matrix ***X*** is the product of two nonnegative matrices ***W*** and ***H***:

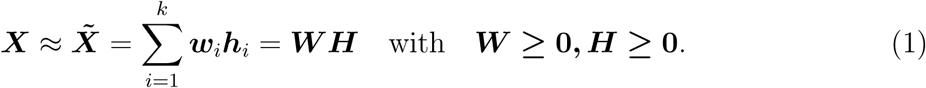

Matrix ***W*** ↓ ℝ*^M↔k^* contains spatial signatures along the pixel domain, while ***H*** ↓ ℝ*^k↔N^* contains spectral signatures along the *m/z* -domain, with the *i*-th NMF-component combining the *i*-th spatial signature with the *i*-th spectral signature. After applying NMF, spatial correlation is captured by ***H*** in spectral signatures grouping ion species that report a similar spatial pattern (captured in ***W***). A detailed treatment on NMF decomposition is provided in supplementary section S2.1. Note that the NMF-supplied lower-dimensional space is not the final IsoSpace-supplied lower-dimensional space.

### 2.2 IsoSpace stage 2 - Spectral filtering of spatially similar ions

Stage 2 seeks to filter spatially similar ion images down to a subset of potential isotope candidate images. Post-NMF, spatial similarity lies encoded in the spectral signature rows of ***H***, and criterion (ii) for isotope candidates is met without requiring prior knowledge or human curation. These rows serve as inputs to stage 2, where we ensure that grouped ion species additionally satisfy the spectral criterion for isotopes, namely that their peaks lie at a *m/z* -distance of 1 *± ɛ* from each other. To incorporate this spectral cue, we construct an *m/z* -pattern matrix to filter the NMF-derived groupings, yielding a refined subset of potential isotope images that are consistent both spatially and spectrally. Each component in the basis matrix ***H*** is processed separately during stage 2.

#### 2.2.1 Construction of ***m/z***-based pattern matrix

To discover spectral patterns-of-interest, we construct an *m/z* -pattern matrix that describes connections between nodes that represent *m/z* -peaks in the IMS data. This matrix facilitates a search for relationships between peaks, using sought-after *m/z* -differences as parameters.

In this paper, the *m/z* -pattern matrix is used to hunt for potential isotope relationships, and is constructed as follows:

***Step 1:*** We compute a *m/z* -distance matrix ***D*** ↓ ℝ*^M↔M^* from the *m/z* -values of all *M* peaks in the IMS dataset. Given a vector of *m/z* -values of size *M* (*e.g.*, *M* = 879 in the mouse pup dataset, *M* = 2199 in the human kidney dataset, and *M* = 2611 in the rat brain dataset), the distance between each possible pair of *m/z* -values, (*m/z*)*_i_* and (*m/z*)*_j_*, is calculated using

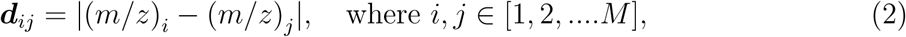

and stored in the *i*^th^ row and *j*^th^ column of matrix ***D***. The absolute *m/z* -differences ensure that the resulting distance matrix ***D*** is both symmetric and non-negative.

***Step 2:*** We identify connections-of-interest between peaks by filtering ***D*** for *m/z* -distances-of-interest, using a tolerance around the desired *m/z* -distance to accommodate for empirical deviations. Focusing on isotope candidates, the *m/z* -distance-of-interest is specified as 1 *±* ɛ.

We construct a primary connection matrix ***C*** ↓ ℝ*^M↔M^*, where each non-zero entry ***c****_ij_* denotes the presence of an edge between nodes (*m/z*)*_i_* and (*m/z*)*_j_*, *i.e.*, their *m/z* -distance ***d****_ij_* falls within the desired envelope:

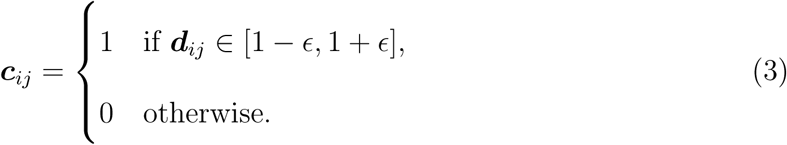

The binary matrix ***C*** indicates which peak pairings fit the *m/z* -distance-of-interest, capturing potential isotope connections for each peak.

***Step 3:*** Once ***C*** reports peak-to-peak connections-of-interest, an iterative exploration, inspired by depth-first search techniques, sequentially establishes potential isotopic ladders for each *m/z* -peak. This step expands the discovery of two-peak patterns to greater-than-two-peak patterns. The process starts with an initial peak (*m/z*)*_i_*. For each such peak, we seek the next peak (*m/z*)*_j_* located within 1 *±* ɛ Da. Since our datasets are MALDI-TOF data, we reasonably assume peaks are singly-charged, but the algorithm can be easily adjusted to accommodate multiple charging as well. The iterative exploration method constructs a pattern matrix ***P*** ↓ ℝ*^M↔M^* to keep track of discovered potential isotopic ladders among the given *M m/z* -values. When multiple peaks satisfy the 1 *± ɛ* distance criterion, we record all such *m/z* -peaks, but continue the search for the next candidate from the first occurrence. Specifically, after identifying (*m/z*)*_j_* relative to (*m/z*)*_i_*, the search progresses from (*m/z*)*_j_* to the subsequent peak (*m/z*)*_k_* spaced at a spectral distance of 1 *±* ɛ, continuously updating ***P*** to capture the emerging isotopic sequences. This process leads to dynamic updates of *m/z* - pattern matrix ***P***, while searching. The search terminates when no further peaks meet the proximity criterion relative to the current reference peak. Matrix ***P*** is updated as follows:

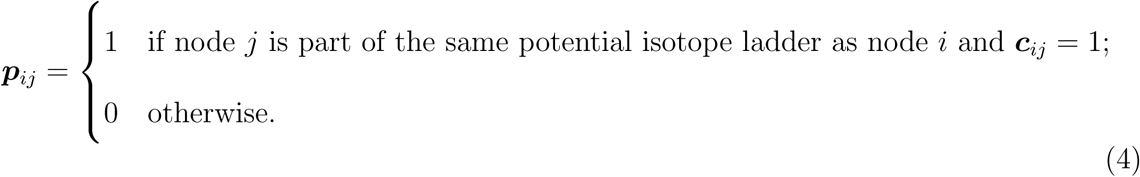

In matrix ***P***, the *i*^th^ column corresponds to the *i*^th^ *m/z* -value out of *M* peaks. Within the *i*^th^ column, each row corresponds to another peak. The rows that are carrying a nonzero value indicate *m/z* -values that are at a distance of 1 Da (allowing for an 2*ɛ* window) or at multiples of this distance from the *i*^th^ peak. For example, if (*m/z*)*_i_* is 1*±*ɛ away from (*m/z*)*_j_*, and (*m/z*)*_j_* is 1 *±* ɛ away from (*m/z_k_*), the (*m/z*)*_i_* column will contain nonzero values in the (*m/z*)*_i_*, (*m/z*)*_j_*, and (*m/z*)*_k_* rows. This results in an asymmetric matrix design that aids in detecting potential isotope candidate patterns by serving as a dataset-specific database of potential hits. Details on this process, using a subset of the mouse pup dataset’s peaks as an example, are provided in supplementary section S2.2.

#### 2.2.2 Detection of isotope candidate patterns by combining the ***m/z***-pattern matrix and NMF-components

After grouping spatially similar peaks in ***H*** and obtaining spectral connections-of-interest in ***P***, each row of ***H*** is analyzed for peak sets whose members share both spatial co-expression and isotope candidate-reporting spectral distances. The procedure to systematically discover such groups of isotope candidates involves:

1. **Peak selection**: In the A^th^ component (row) of ***H***, select the peak with the highest component intensity.
2. **Connection discovery**: Retrieve the selected peak’s column in ***P*** . The column’s nonzero rows indicate peaks successively spaced at 1 *±* ɛ, highlighting how the selected peak fits into a sequential 1 *±* ɛ Da progression, *i.e.*, a potential isotopic ladder.
3. **Pattern extraction**: Using the A^th^ component vector in ***H***, keep the entries that match active rows in ***P*** ’s column and set the others to zero. This pattern vector, designated “pattern A-1”, effectively reports *m/z* -values or peaks that demonstrate both spatial co-expression and isotope candidate-reporting spectral distances. The found isotope candidate pattern is stored as a new row in matrix ̃H ɛ ℝ*^Q×N^*, where *Q* is the number of extracted patterns, and it functions as a new latent axis for IsoSpace’s chemistry-informed DR.
4. **Repetition avoidance**: In the A^th^ component vector, set entries corresponding to peaks active in the extracted pattern to zero, to prevent re-use of the same peaks in subsequent pattern searches within the same component.
5. **Intra-component iteration**: Repeat steps 1-4, selecting the next highest peak that still exceeds a preset component intensity threshold ” and extracting the next isotope candidate pattern. This iterative process continues until no peaks exceeding ” remain.
6. **Inter-component iteration**: Repeat steps 1-5 for each component in ***H***, systematically cataloging for the entire dataset spatially correlated patterns of isotope candidates as individual rows in ̃H.

At the end of this procedure, matrix ̃H contains Q rows and effectively decomposes empirically measured mass spectra into a combination of its *Q* components. IsoSpace takes an IMS dataset where every pixel is a point in a *M* -dimensional space and delivers a lower-dimensional representation where every pixel is a point in a *Q*-dimensional space. This is similar to DR-methods like PCA and NMF, albeit that IsoSpace’s components are directly interpretable as potential isotopic families of peaks, *i.e.*, molecular species. As a bonus, each of the Q components is an empirically extracted set of peaks that are potential isotopes of each other.

## 3 Results and Discussion

Our algorithm is tested on the described IMS datasets, with five distinct case studies exploring the quality of results and hyperparameter impact.

### 3.1 Impact of the ***m/z***-uncertainty parameter ɛ

The first case study explores the impact of the ɛ-parameter, which sets the *m/z* -uncertainty permitted around the spectral distance criterion for potential isotopic patterns. We investigate how changes in the ɛ-parameter a!ect the quantity and lengths of the detected isotope candidate patterns. Specifically, we examine ɛ-values of *m/z* 0.03, 0.01, 0.009, 0.007, 0.005, and 0.003, and we evaluate the relationship between the spectral tolerance and characteristics of the extracted patterns. This study analyses, in the context of IMS data analysis, IsoSpace’s isotope candidate pattern detection and its sensitivity to the ɛ-parameter.

First, we explore how *m/z* -uncertainty, and thus mass resolution, relate to the numberof found isotope candidate patterns. Figure 3a presents, for the mouse pup dataset, the cumulative lengths and occurrences of found isotope candidate patterns as a function of *ɛ*. Our findings indicate that short patterns of two or three peaks tend to dominate the results. They also show that increasing *ɛ* from narrower (little *m/z* -uncertainty, *i.e.*, high mass resolution) to wider *m/z* -windows (more *m/z* -uncertainty, *i.e.*, lower mass resolution), allows for a lower number but longer patterns to be found. This is potentially because distinct short patterns in a fine-mass-resolution regime get connected into one long pattern when a more forgiving *m/z* -precision is allowed. For the mouse pup dataset, an ɛ-value of 0.009 appears optimal, as it enables short patterns to merge into longer isotopic ladders without substantially reducing the overall count. In contrast, larger ɛ-values such as 0.01 and 0.03 yield markedly fewer patterns and introduce unrealistically long ladders.

**Figure 3:**
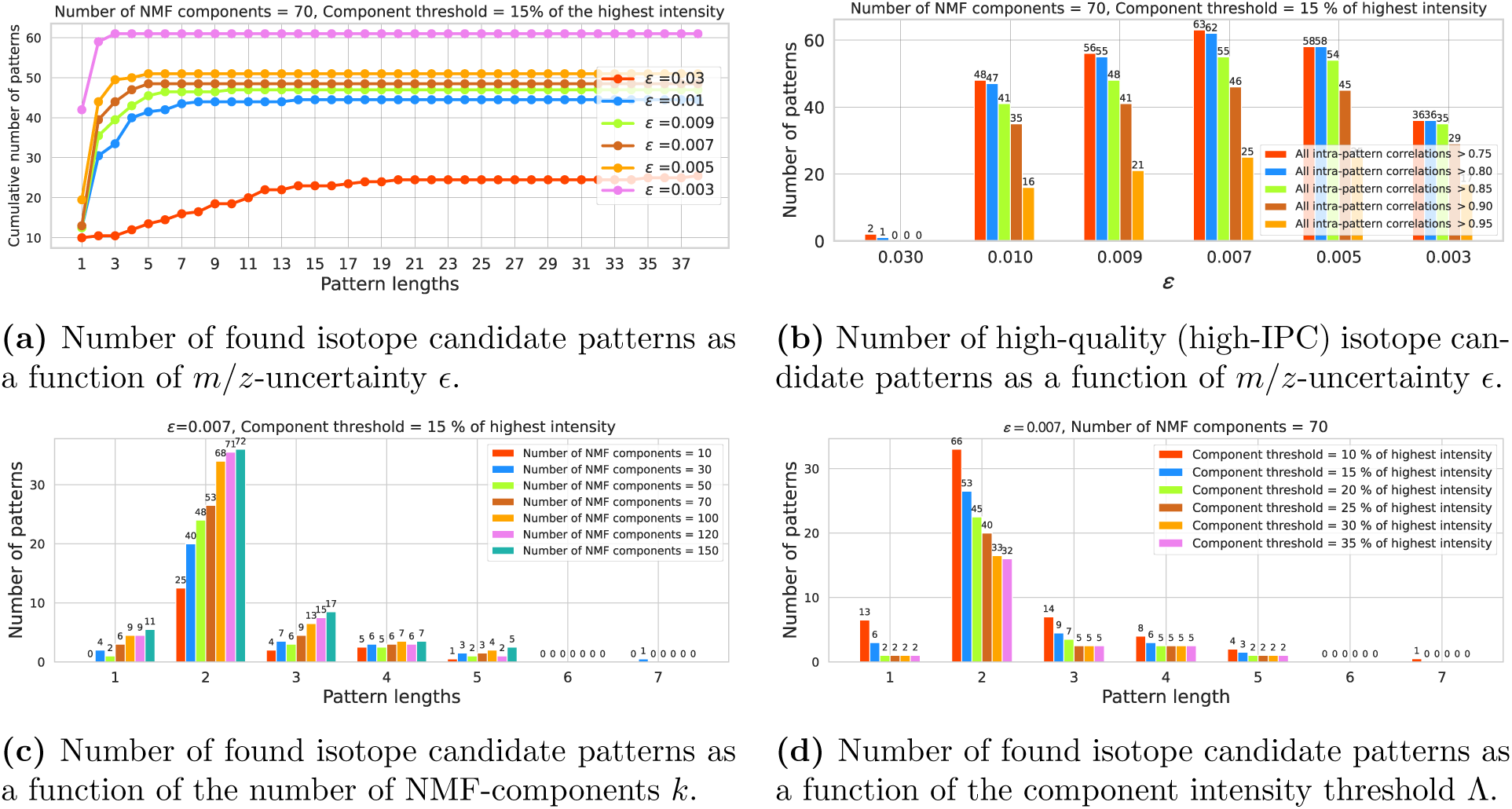
Comparative analysis: (a) Cumulative counts of found isotope candidate patterns and their respective pattern lengths (*i.e.*, number of peaks) in function of ɛ. These results use *k* = 70 NMF-components and a component threshold ” = 15% of the highest component intensity. (b) Number of isotope candidate patterns with IPCs exceeding the given quality threshold, in function of ɛ. (c) Counts of found isotope candidate patterns and their respective pattern lengths in function of the number of NMF-components *k* used in Stage 1. (d) Counts of found isotope candidate patterns and their respective pattern lengths in function of the component intensity threshold ” (with ɛ = 0.007).

Across all ɛ-values, a decline in the total number of patterns coincides with an increase in pattern lengths, illustrating a saturation trend. At the broadest *m/z* -uncertainty allowance (ɛ = 0.03), only a small number of patterns are detected in total, yet they span a wide range of lengths (up to 38 peaks). This indicates that many long but sparse ladders can be formed under loose *m/z* -distance constraints. In contrast, for narrower allowances (ɛ ≃ 0.01), the cumulative number of patterns is substantially higher, but the ladders saturate at shorter lengths (typically 3-5 peaks). This highlights a trade-o!: broader *m/z* -windows capture longer but less reliable patterns, whereas narrower windows recover more isotopically specific ladders of limited length. Moreover, with the narrowest *m/z* -allowance (ɛ = 0.003), patterns of two peaks dominate, failing to find another peak within a 1 *±* 0.003 spectral distance of the first two. Patterns that feature only a single *m/z* -peak are designated as singletons, indicating the absence of isotopic connections to other *m/z* -values for the given ɛ. These observations underscore that a narrower ɛ results in more *m/z* -specific patterns found, but also the elimination of (sometimes many) less *m/z* -precise patterns. As ɛ is allowed to became wider, the found isotope candidate patterns become looser in terms of *m/z* and longer, but sometimes unrealistically long. Also, there seems to be an ɛ lower band, below which isotope relationships go undetected, and a critical threshold above which isotopic relationships become so loosely defined as to become indiscernible.

Besides assessing the number and length of found isotope candidate patterns, it is also crucial to evaluate their quality. To accomplish that, we check the overall spatial correlation among the ions that a pattern pulls together. We define intra-pattern correlation (IPC) as a measure to gauge that spatial coherence. IPC calculates the pairwise correlation among all ion images in a found pattern. If the peaks in the pattern are genuine isotopes of each other, their spatial distributions should be nearly identical, save for absolute intensity-related differences and noise. Thus, in an isotopic pattern, we expect strong spatial correlation between the ion images and a high IPC. By setting an IPC-threshold, we can determine whether all pairwise correlations within a found pattern exceed this threshold. We label patterns that meet this criterion as “homogeneous”, indicating strong spatial correlation among all constituent ion images and providing a qualitative assessment of the isotope candidate patterns’ coherence. Figure 3b shows IPCs for different ɛ-values in the mouse pup dataset, using various quality thresholds. The plot counts the number of patterns (excluding singletons) whose IPC exceeds the specified thresholds. The number of these homogeneous patterns, which have an increased probability of reporting genuine isotopes, rises as ɛ-values decrease and transition from wider to narrower *m/z* -windows. However, at ɛ-values smaller than 0.007, the count declines again, most probably because ɛ becomes too narrow and restrictive. Overall, Figure 3b suggests that as ɛ becomes more stringent, the quality of the found isotope candidate patterns goes up. However, as ɛ is set narrower and narrower, at a certain moment, ɛ has become so narrow and exacting that no more isotopes are found. This trend highlights the nuanced relationship between ɛ-values and the spatial coherence of found patterns, underscoring the need for a balance between allowance for *m/z* -uncertainty and result quality. Note, at the widest and most forgiving ɛ, the number of patterns is quite high (Fig. 3a), while the number of high-quality patterns drops to nearly 0 (Fig. 3b). Furthermore, as we raise the IPC-threshold to demand higher-quality patterns, the number of detected homogeneous patterns declines overall, but it seems relatively more so for broader *ɛ*-windows. This could suggest increased challenge in finding high-quality isotope candidate patterns under worse mass resolution conditions.

The observations from Figures 3a and 3b can help users make informed choices for ɛ, by tailoring it to characteristics of their dataset and to the requirements of their application. For example, is it better for the user’s study to employ a narrower ɛ-window to obtain strengthened IPC and higher-quality isotope candidate patterns, albeit at the cost of reduced pattern lengths? Or is it more useful to select an ɛ-value that compromises between IPC and pattern length, taking into account the empirical mass resolution of the dataset? For the mouse pup dataset, *ɛ* = 0.007 seems an optimal choice, balancing the *m/z* -precision of isotope candidate pattern detection with the number of found patterns and their quality. Specifically, with 879 measured *m/z* -peaks, an *m/z* -uncertainty of *ɛ* = *±*0.007, and *k* = 70 NMF-components, IsoSpace discovered 53 patterns of two isotope candidates, 9 patterns of three isotope candidates, 6 patterns of four isotope candidates, and 3 long patterns of merged isotope candidate ladders. The patterns found account for 172 of 879 measured *m/z* -peaks, suggesting that 172 peaks could be replaced by their 71 monoisotopic peaks in downstream analyses. This means that IsoSpace alone could reduce the dimensionality of this dataset by 11.5% with minimal loss of chemical information. Replacing isotopic relatives by a single peak representing the molecular species would not only aid human interpretation and storage requirements but would also reduce collinearity issues in subsequent computational analysis. The results of similar experiments on the rat brain and HuBMAP datasets are provided in the supplementary section S3.2.

### 3.2 Impact of the number of NMF-components

The second case study investigates how the number of NMF-components, *k*, impacts detection of isotope candidate patterns. Figure 3c shows results for *k*-values 10, 30, 50, 70, 100, 120, and 150. Patterns of two peaks are found most frequently, with their occurrence increasing as *k* rises. For longer pattern lengths, the number of patterns found tends to decrease for all *k* and they seem to become less sensitive to the particular choice of *k*. For the mouse pup dataset, increasing *k* in the first stage of IsoSpace results in an increase of the number of found isotope candidate patterns of length two and three. However, for finding isotope candidate ladders of four or more peaks, the number of components *k* does not seem to be of much influence. In the mouse pup dataset, a *k*-value of 70 emerged as a good setting. Overall, the choice of *k* seems like it should probably be primarily driven by whether the finding of short isotopic ladders is desirable. Similar results on the other datasets can be found in supplementary section S3.3.

### 3.3 Impact of the component intensity threshold ”

The third case study examines the effect of the component intensity threshold ” on finding isotope candidate patterns. Figure 3c shows the results for ”-values of 10, 15, 20, 25, 30, and 35. There is a predominance of length-two patterns, with a gradual decrease in detected patterns as ” increases. A lower ” yields a higher number of patterns detected, for all pattern lengths. As ” increases, fewer patterns are discerned. Longer isotope candidate patterns seem less a!ected by a rising ”, while short two-peak patterns are found less often as ” goes up. This underscores ”’s role for modulating the detectability and diversity of patterns. For the mouse pup dataset, we have considered a ”-setting of 15% of the highest component intensity for all experiments. Overall, a lower ” enhances pattern detection, albeit potentially including duplicates from different NMF-components. A higher ” restricts the sensitivity of the isotope candidate pattern detection, but can relieve some of the computational resources needed for a detection run. These insights facilitate a data-driven approach to ”-selection, optimizing for discovery sensitivity under given resources.

### 3.4 Visualizing isotope candidates in function of ***ɛ***

The fourth case study provides a visual examination of isotope candidate patterns through their ion images. We vary ɛ to see how different mass resolution scenarios impact individual patterns. Figure 4 shows two specific patterns that were detected in an IsoSpace run on the mouse pup dataset. In Figures 4a and 4c, at intermediate *ɛ*-values such as 0.007 or 0.009, genuine isotopes of *m/z* 769.578 seem to be found in *m/z* 770.582 and 771.587. At a very narrow ɛ-value, such as 0.003, the mass error of the measurement seems larger than the allowed uncertainty and none of the isotopes are found and connected. At a very wide ɛ-value, such as 0.030, the allowed uncertainty facilitates connection to additional peaks: *m/z* 772.602, 773.609, 774.616, and 775.619. While from the spectral perspective of Figure 4a these peaks look like they could be isotopes, the ion images in Figure 4c show that they are, in fact, false positives that got spuriously connected to *m/z* 769.578 as the uncertainty ɛ was opened up too wide. These observations show that ɛ-values from 0.005 to 0.010 successfully capture isotopes within 1 *±* ɛ distances from each other while also showing similar tissue distributions in their ion images. In Figures 4b and 4d, at a very narrow ɛ-value of 0.003, none of the isotopes are found and connected, while at an *ɛ* of 0.005 a set of three isotope candidates are linked to each other: *m/z* 713.465, 714.470, and 715.468. Furthermore, at *ɛ*-values 0.007 through 0.010, *m/z* 716.474 is added as an isotope candidate as well. From the spectral perspective in Figure 4b, *m/z* 716.474 looks like it could be a genuine member of the isotopic ladder starting at *m/z* 713.465. However, the ion images in Figure 4d show that it is actually a false positive, incorrectly connected due to wide *ɛ*-uncertainty allowance.

**Figure 4:**
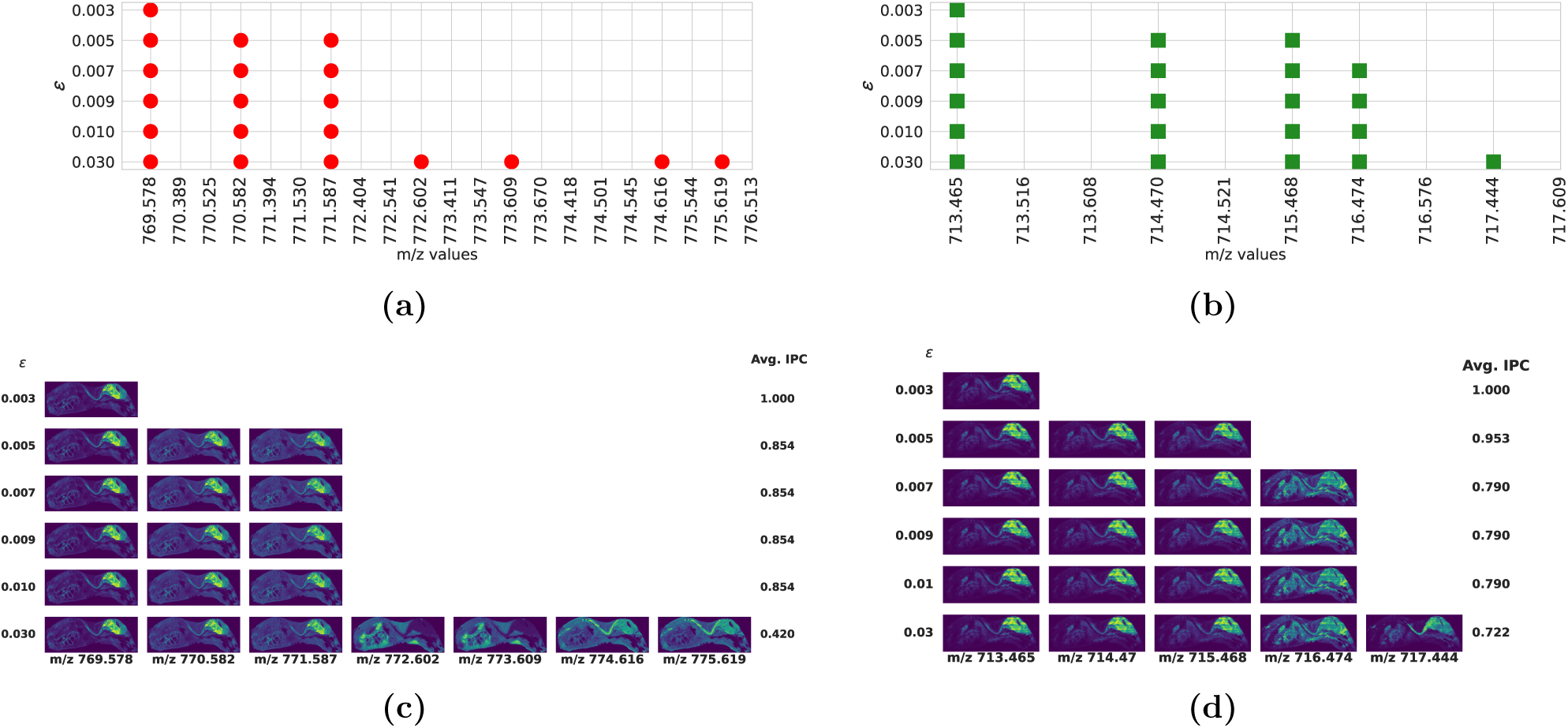
Spectral and spatial views of two isotope candidate patterns found by IsoSpace, consid-ering different levels of ɛ *m/z* -uncertainty. (a) Isotope candidate pattern starting at *m/z* 769.578. A red dot means that a *m/z* -value is part of the pattern at the corresponding ɛ-value. (b) Isotope candidate pattern starting at *m/z* 713.465. A green square means that a *m/z* -value is part of the pattern at the corresponding ɛ-value. (c) Ion images for the isotope candidate pattern starting at *m/z* 769.578. (d) Ion images for the isotope candidate pattern starting at *m/z* 713.465. Note the IPCs listed for each combination of ion images, grading their pattern quality at a particular ɛ.

While visual assessment from ion images works, it does not scale to large numbers of patterns. Instead we will use IPC to gauge the pattern match numerically. For example, from ɛ-value 0.007 onwards to 0.030, the set of peaks does not exhibit a high IPC. This suggests that non-isotopes have entered the pattern or that the added ion image is a mixture of a genuine isotope with another ion species. At an *ɛ* of 0.005, the pattern consists solely of highly correlated ion images, which is reflected in the high IPC of 0.953. This analysis demonstrates the delicate balance between *ɛ* and the quality and fidelity of the detected isotope candidate patterns. It highlights the importance of choosing an appropriate *ɛ* that still captures isotope-like relationships while also maintaining high spatial correlation between those isotope candidates’ images. Overall, narrower *ɛ*-windows struggle to detect isotopes due to too strict spectral distance constraints. Conversely, as the *ɛ*-window widens, additional *m/z* -peaks are recognized as potential isotopes, albeit typically with a lower average intra-pattern correlation (IPC) as a result.

### 3.5 Isotope candidate patterns across different IMS datasets

The fifth case study investigates isotope candidate patterns in the rat brain and human kidney (HuBMAP) datasets. Figure 5 shows two isotope candidate patterns retrieved by IsoSpace from the rat brain dataset. The spectral plots on the left (Figs.5a and 5c) highlight the peaks present in the NMF-component of ***H*** in red, the component intensity threshold ”in blue, and the found isotope candidate pattern in cyan. The corresponding ion images are displayed on the right (Figs.5b and 5d), showcasing the spatial distributions of the selected peaks. Isotope candidate pattern 5-56 (Figure 5a) extends from *m/z* 5436.197 to 5443.196, with peaks separated by 1 *± ɛ* where *ɛ* = 0.020 and *k* = 70. The *m/z* -value preceding this pattern is distanced 27.689 Da away, and the subsequent *m/z* -value at 5451.004 is 7.808 Da away, correctly excluding them from the pattern. The strong spatial similarity between the ion images of pattern 5-56 (Figure 5b) suggests that the found isotope candidates are probably legitimate isotopes.

**Figure 5:**
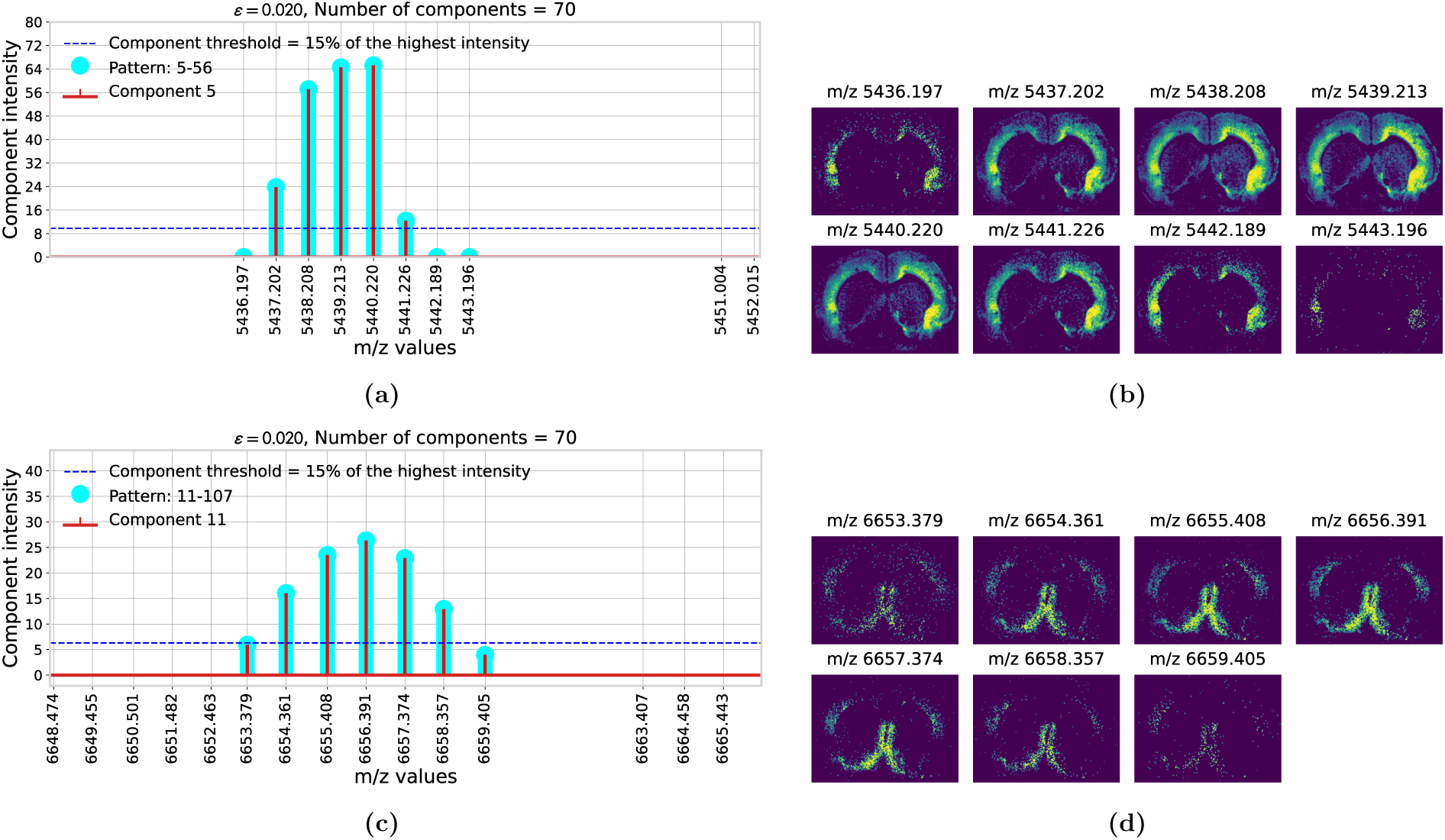
Isotope candidate patterns 5-56 and 11-107, detected in the rat brain dataset. (a)(c) Spectral plots highlight in red the peaks present in the NMF-component of ***H***, in blue the com-ponent intensity threshold ”, and in cyan the found isotope candidate pattern. (b)(c) The corre-sponding ion images show the spatial distributions of the selected peaks.

Although perhaps not immediately apparent, in isotope candidate pattern 11-107 (Figure 5c), not all *m/z* -peaks adhere to the 1*±*0.020 spacing criterion. For instance, the spacing between *m/z* 6654.361 and 6655.408 is 1.047, and between *m/z* 6658.357 and 6659.405 is 1.048. Both distances exceed the 1 *±* 0.020 envelope. Thus, without further modification, IsoSpace would eliminate such *m/z* -peaks from participating in this pattern. In these situations, where the detection of an *m/z* -peak may be hindered by its low signal intensity or by falling just outside the preset *m/z* -window, it can be challenging to detect an entire isotope candidate sequence without interruptions. Hence, we introduced into IsoSpace the option to have “missing values” in the *m/z* -pattern matrix. This allows the algorithm to accommodate for coincidental dropouts of *m/z* -peaks. IsoSpace addresses this by not only searching for *m/z* -distances of 1 *± ɛ*, but also spacings of 2 *±* 2*ɛ*. This 2*ɛ*-adjustment is designed to account for one missing peak in a growing pattern. Specifically, IsoSpace proactively seeks the next isotope candidate peak at a distance of 1 *± ɛ*. Failing that, it extends the search to 2 *±* 2*ɛ*, maintaining sequence integrity and o!ering robustness against occasional gaps.

Reconsidering pattern 11-107, the number of missing values is set to 1, with *ɛ* = 0.020 and *k* = 70. The analysis begins with the first peak at *m/z* 6653.379. This peak successfully connects with the second peak at *m/z* 6654.361, as their distance fits within the *ɛ*-window. However, from *m/z* 6654.361, the algorithm does not find the next peak within the 1 *± ɛ* envelope, leading to a temporary end of the isotope candidate sequence. Given the new allowance for one missing peak, the algorithm then links the second peak, *m/z* 6654.361, to the fourth peak at *m/z* 6656.391 with their distance falling within the 2 *±* 2*ɛ* range. Subsequently, from *m/z* 6656.391, the algorithm searches in both forward and backward directions for the next peak within the 1 *± ɛ* range. This leads to the detection of the third peak, *m/z* 6655.408, and its incorporation into the isotope candidate sequence after all. Additional examples demonstrating the utility of the missing value hyperparameter and details on the construction of the corresponding *m/z* -pattern matrix are available in the supplementary information. These results highlight IsoSpace’s effectiveness in ensuring isotopic sequence coverage in the face of missing data. Figure 6 shows two isotope candidate patterns retrieved by IsoSpace from the human kidney dataset. Figure 6a illustrates one such isotope candidate ladder, from *m/z* 453.200 to 455.212, using *ɛ* = 0.007 and *k* = 70. The *m/z* -value immediately preceding this sequence, *m/z* 452.192, is 1.008 Da away. This distance exceeds the 1 *±* 0.007 envelope and the peak is therefore excluded under this narrow *ɛ*-window. The corresponding ion images (Figure 6b) demonstrate strong spatial similarity among the peaks, suggesting that the found isotope candidates are probably legitimate isotopes. A second pattern is shown in Figures 6c and 6d, where the inclusion of closely spaced neighboring peaks is prevented by the use of the narrower *m/z* -window *ɛ* = 0.007, enabling the extraction of the true isotopic pattern.

**Figure 6:**
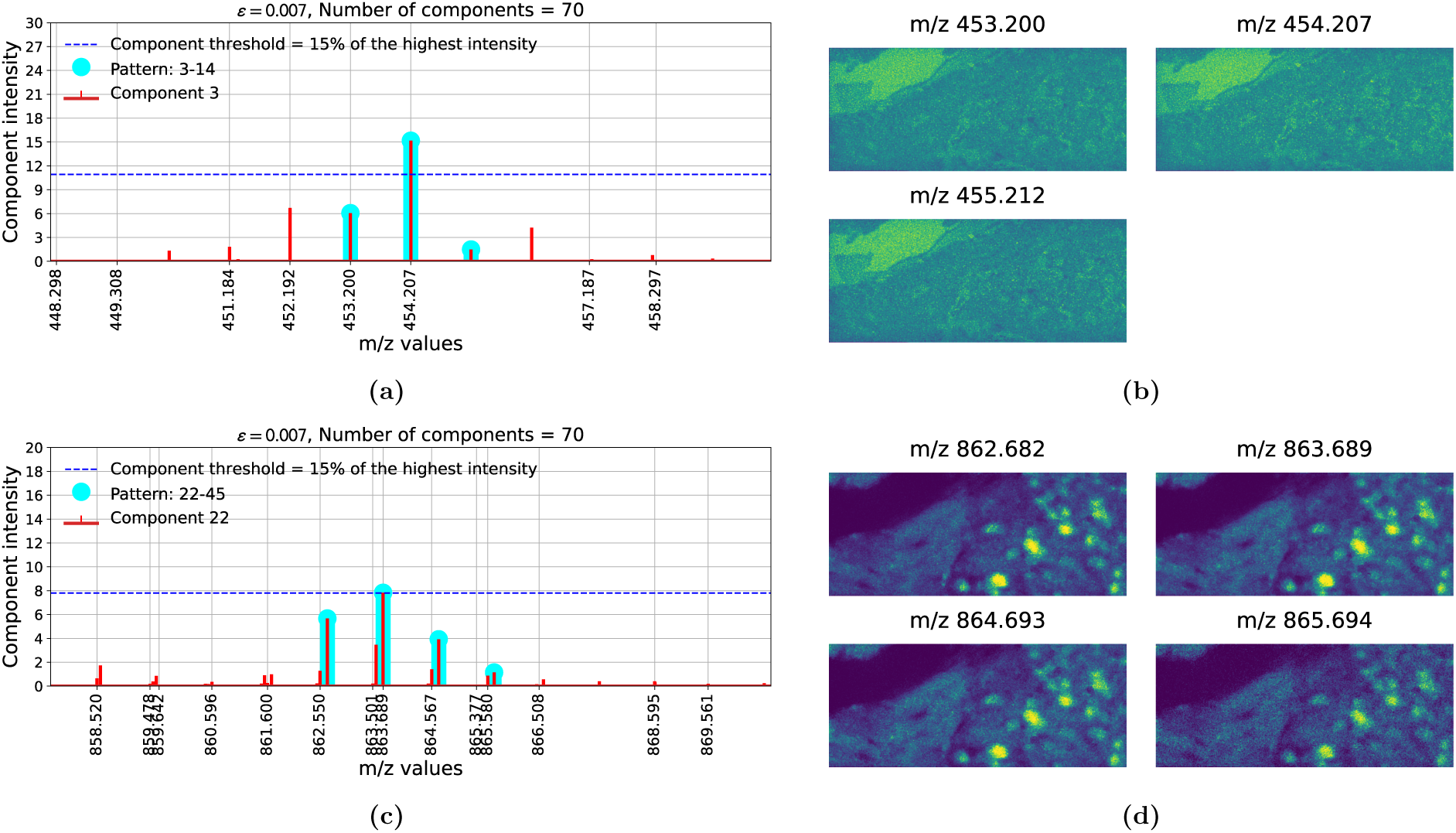
Isotope candidate patterns 3-14 and 22-45, detected in the human kidney dataset. (a)(c) Spectral plots highlight in red the peaks present in the NMF-component of ***H***, in blue the component intensity threshold ”, and in cyan the found isotope candidate pattern. (b)(c) The corresponding ion images show the spatial distributions of the selected peaks.

Finally, in Figure 5a, the pattern’s tail end features three peaks with notably lower component intensity compared to other peaks. Their ion images display diminished intensity, leading to a lower IPC. Although their spatial distributions remain consistent with peaks of higher intensity, the lower abundance of these isotope candidates can interfere with the pattern’s overall quality assessment. To refine our analysis, we introduce an isotope candidate abundance threshold #. This threshold is defined as a percentage of the highest component intensity, and facilitates exclusion of lower-intensity peaks to enhance IsoSpace’s analytical precision. This thresholding approach not only ensures that the analysis focuses on the most significant and abundant isotope-like signals, but it also improves data interpretability and computational e$ciency. By filtering out less abundant isotope candidates, our methodology streamlines the identification of pertinent isotopic patterns within complex IMS datasets, as illustrated by an example in the supplementary information.

## 4 Conclusion

IsoSpace seeks to cast an IMS measurement from its original ion species-reporting space to a molecular species-reporting space. In the input, the measurement space, each feature or dimension reports a different ion species. In the output, a lower-dimensional latent space, each feature or dimension reports a different molecular species, after combining related isotopes into a single reporting feature. This chemistry-informed DR for IMS data represents a substantial advancement in the field, offering a unique approach to disentangling a complex spatially-resolved mixture of isotopic peaks without prior assumptions about molecule classes. The lower-dimensional IMS dataset representation it provides, serves the classical purposes of DR, but sacrifices some compression to maintain better chemical interpretability and to suggest potential isotopic relationships among measured ion species. Across diverse case studies, IsoSpace enhances data interpretability, but also performs DR tasks such as data compression and minimizing collinearity prior to follow-up computational analyses. IsoSpace’s ability to handle missing peaks in isotopic patterns, detailed control through hyperparameters, and strong intra-pattern correlation results despite noise make it essential for detecting spatially-consistent isotope candidates. This work highlights what an automated, data-driven approach for chemistry-informed DR and isotope candidate detection can bring to IMS research. By filtering out redundant information and focusing on the most chemically relevant features, researchers can more easily interpret the data, identify compounds, and understand the chemical composition of the samples under investigation.

## Acknowledgement

Research reported in this publication was supported by the National Institutes of Health (NIH)’s Common Fund, National Institute Of Diabetes And Digestive And Kidney Diseases (NIDDK), and the O$ce Of The Director (OD) under Award Numbers U54DK120058, U54DK134302, and U01DK133766 (J.M.S. and R.V.), by NIH’s Common Fund, National Eye Institute, and the O$ce Of The Director (OD) under Award Number U54EY032442 (J.M.S. and R.V.), by NIH’s National Institute Of Allergy And Infectious Diseases (NIAID) under Award Numbers R01AI138581 and R01AI145992 (J.M.S. and R.V.), by NIH’s National Institute On Aging (NIA) under Award Number R01AG078803 (J.M.S. and R.V.), by NIH’s National Cancer Institute (NCI) under Award Number U01CA294527 (J.M.S. and R.V.), and by the National Science Foundation Major Research Instrument Program CBET – 1828299 (J.M.S.). The research was furthermore made possible in part by grant numbers 2021-240339 and 2022-309518 (L.G.M. and R.V.) from the Chan Zuckerberg Initiative DAF, an advised fund of Silicon Valley Community Foundation. The content is solely the responsibility of the authors and does not necessarily represent the o$cial views of the National Institutes of Health.

## Footnotes

^1^https://portal.hubmapconsortium.org/browse/dataset/271efc13304fbec50b1fe3c111ceea3e

^2^doi:10.35079/HBM437.BGCD.226 VAN0042-RK-1-31-IMS lipids neg multilayer.ome

